# AIRRSHIP: simulating human B cell receptor repertoire sequences

**DOI:** 10.1101/2022.12.20.521228

**Authors:** Catherine Sutherland, Graeme J M Cowan

## Abstract

Adaptive Immune Receptor Repertoire Sequencing is a rapidly developing field that has advanced understanding of the role of the adaptive immune system in health and disease. Numerous tools have been developed to analyse the complex data produced by this technique but work to compare their accuracy and reliability has been limited. Thorough, systematic assessment of their performance is dependent on the ability to produce high quality simulated datasets with known ground truth. We have developed AIRRSHIP, a flexible and fast Python package that produces synthetic human B cell receptor sequences. AIRRSHIP uses a comprehensive set of reference data to replicate key mechanisms in the immunoglobulin recombination process, with a particular focus on junctional complexity. Repertoires generated by AIRRSHIP are highly similar to published data and all steps in the sequence generation process are recorded. These data can be used to not only determine the accuracy of repertoire analysis tools but can also, by tuning of the large number of user-controllable parameters, give insight into factors that contribute to inaccuracies in results.

**Availability and Implementation:** AIRRSHIP is implemented in Python. It is available via https://github.com/Cowanlab/airrship and on PyPI at https://pypi.org/project/airrship/. Documentation can be found at https://airrship.readthedocs.io.

## Introduction

Adaptive Immune Receptor Repertoire Sequencing (AIRR-seq) uses high throughput sequencing to characterise the state and dynamics of B cell receptor (BCR) repertoires and is of increasing use in understanding the molecular basis of immunity and autoimmunity (Yaari and Kleinstein, 2015; Zheng *et al.*, 2022). BCR repertoire data comprises sequences of the variable regions of BCRs, which are rearranged and hypermutated during B cell maturation. Key to interpretation of this data is the determination of parental VDJ segments for each receptor, as well as identification of nucleotide insertions and deletions, positions of segment junctions, and sequence mutations. Numerous tools have been developed to enable these analyses, with the most commonly used being IgBLAST, IMGT/HighV-QUEST and MiXCR (Brochet *et.al*, 2008; Ye *et al.*, 2013; Bolotin *et al.*, 2015). Benchmarking of such tools has been limited but work published to date indicates that their outputs can differ, potentially impacting the conclusions drawn from data (Smakaj *et al.*, 2019).

The ability to assess the absolute accuracy of antibody repertoire analysis tools is limited by the lack of experimental datasets where details of all recombination processes are known. Simulation of rearranged BCR data offers an alternative solution for benchmarking where this ground-truth is recorded. Several methods for simulating BCR data have been published (Ralph and Matsen, 2016; Yang *et al.*,2021; Weber *et al.*, 2020; Safonova *et al.*, 2015; Marcou *et al.*, 2018; Han *et al.*, 2022; Yermanos *et al.*, 2017). However, the usefulness of these tools for benchmarking studies can be limited by their requirements for complex dependencies, non-standard output file formats and/or generation of repertoires that are dissimilar to real experimental data. Therefore, we introduce AIRRSHIP (Adaptive Immune Receptor Repertoire Simulation of Human Immunoglobulin Production), a tool for simulating human BCR sequences, which accurately replicates features of experimental data whilst being fast, flexible, and easy to use.

## Implementation

AIRRSHIP is implemented in Python and has no additional dependencies beyond the standard library. It replicates the processes by which immunoglobulin sequences are formed *in vivo*, with each step informed by comprehensive built-in reference data from experimental datasets (Supplementary Tables 1 & 2). VDJ segments are selected from simulated immunoglobin loci according to observed patterns of usage, trimming of segment ends then occurs and nucleotides are inserted at junctions (Figure 1). Somatic hypermutation is replicated at both the per sequence and per position level, with bases mutated according to both local and wider sequence context. Models vary at the IMGT gene, family or sequence region level and thus should be robust to novel allele discovery (Lefranc, 2007; Gadala-Maria *et al.*, 2015).

**Figure 1:**
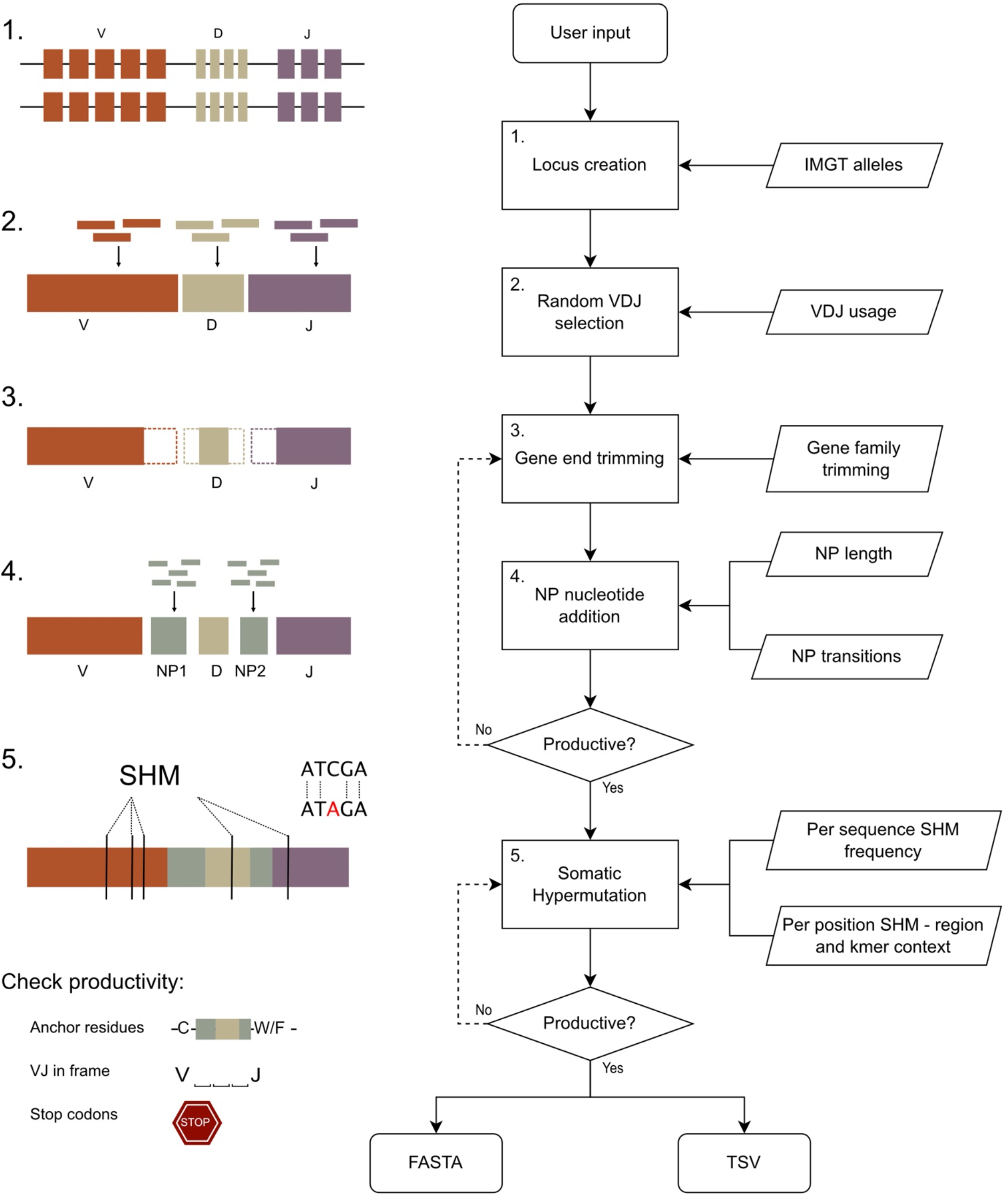
AIRRSHIP simulates human heavy chain BCR sequences, informed by experimental data at each step of the synthetic recombination process. Key parameters such as VDJ usage, gene trimming, junctional insertions and somatic hypermutation can all be modified by the user. Sequences in the FASTA output can then be used as input for tools of interest and results easily compared to the tab separated values (TSV) file which acts as a record of the recombination process for each sequence.

To create repertoires tailored to specific benchmarking scenarios, the user can control patterns of VDJ usage, insertions, and deletions, as well as the rate and distribution of somatic hypermutation. The output is also optimised for benchmarking, consisting of a FASTA file of sequences which can be used directly as input for tools of interest, as well as a tab separated values file, which closely follows the AIRR standards rearrangement schema (Vander Heiden *et al.*, 2018) and facilitates easy reference to known sequence origins.

## Results

We assessed the resemblance to real data of sequences generated by AIRRSHIP and three other simulation tools: partis, immuneSIM and IMPlAntS (Ralph and Matsen, 2016; Weber *et al.*, 2020; Yang *et al.*, 2021). Key summary statistics from these repertoires were compared to those from published BCR data using sumrep (Olson *et al.*, 2019). To prevent favourable bias, experimental data were selected from studies not used for AIRRSHIP input reference (Supplementary Table 3).

Comparison between non-mutated simulated repertoires and IgD/IgM experimental sequences revealed similarly high levels of performance between AIRRSHIP, IMPlAntS and immuneSIM in VDJ usage frequency, with substantial deviation from experimental data in repertoires generated using partis (Supplementary Figure 1). Divergences between real sequences and the AIRRSHIP and IMPlAntS repertoires were similar for lengths of nucleotide deletions and insertions at the junctions, whilst sequences from partis and immuneSIM differed more. However, the composition of nucleotide insertions in AIRRSHIP sequences is much closer to published data due to explicit modelling of nucleotide addition. AIRRSHIP repertoires also most closely resemble experimental data in amino acid usage, GRAVY score, aliphatic index, and Atchley factor scores; all of which are important measures of sequence composition. Similar patterns were observed when comparing simulated repertoires with mutation to experimental IgA/IgG sequences (Supplementary Figure 2).

During affinity maturation, mutations are introduced unevenly across BCR sequences; therefore, we compared the ability of the simulation tools to produce realistic mutation patterns (Supplementary Figure 3). Mutation rates per nucleotide position in the sequence were very closely correlated between AIRRSHIP and experimental sequences (r = 0.97), with a stronger relationship than shown with IMPlAntS (r = 0.95), partis (r = 0.29) and immuneSIM (r = 0.27) repertoires.

Finally, repertoire sequencing datasets may now consist of hundreds of thousands, if not millions, of sequences. Synthetic datasets of similar scale will be required, and these must be able to be produced within a reasonable timeframe. AIRRSHIP can generate 10,000 unique recombinations with mutation in less than 70 seconds. This is faster than, but comparable to, IMPlAntS and partis, whilst immuneSIM is considerably slower (Supplementary Figure 4 & Supplementary Table 4). Sequence generation by AIRRSHIP scales linearly, and one million sequences can be produced in just under two hours with mutation, or in 13 minutes without (Supplementary Table 5).

## Demonstration

To illustrate a potential application of AIRRSHIP, we compared the ability of IMGT/HighV-QUEST and IgBLAST to accurately identify the number of nucleotides inserted at the VD junction (Supplementary Figure 4). Establishing correct gene boundaries within the junction, especially when the D sequence is short, is known to be a difficult task (Yaari and Kleinstein, 2015) and we found that only half of all sequences had the correct insertion length assigned by both tools. To explore what effect gene trimming has on determining insertions, we also tested these tools using sequences where the D gene was not trimmed. In this case, the proportion of sequences with correct assignments rose to almost 90%, confirming D gene trimming as a significant confounder. By further iterating through AIRRSHIP parameters, a greater understanding of limitations in junctional boundary identification and how these might affect downstream analyses could be gained.

## Conclusion

AIRRSHIP simulates BCR repertoires that closely resemble experimental datasets with particularly faithful representation of junctional diversity. It requires few dependencies, runs quickly, and offers broad choices in parameter selection. We have illustrated how AIRRSHIP may be used to explore challenges in AIRR-seq analyses and believe it can be of value in more systematic assessment of the current AIRR-seq tool landscape.

## References

Bolotin, D.A. et al. (2015) MiXCR: software for comprehensive adaptive immunity profiling. Nat. Methods, 12, 380–381.

Brochet, X. et al. (2008) IMGT/V-QUEST: the highly customized and integrated system for IG and TR standardized V-J and V-D-J sequence analysis. Nucleic Acids Res., 36, W503–W508.

Gadala-Maria, D. et al. (2015) Automated analysis of high-throughput B-cell sequencing data reveals a high frequency of novel immunoglobulin V gene segment alleles. Proc. Natl. Acad. Sci. U. S. A., 112, E862–E870.

Han, J. et al. (2022) Echidna: integrated simulations of single-cell immune receptor repertoires and transcriptomes. Bioinforma. Adv., 2, vbac062.

Vander Heiden, J.A. et al. (2018) AIRR Community Standardized Representations for Annotated Immune Repertoires. Front. Immunol., 9, 2206.

Lefranc, M.P. (2007) WHO-IUIS nomenclature subcommittee for immunoglobulins and T cell receptors report. Immunogenetics, 59, 899–902.

Marcou, Q. et al. (2018) High-throughput immune repertoire analysis with IGoR. Nat. Commun., 9.

Olson, B.J. et al. (2019) sumrep: A Summary Statistic Framework for Immune Receptor Repertoire Comparison and Model Validation. Front. Immunol., 10, 2533.

Ralph, D.K. and Matsen, F.A. (2016) Consistency of VDJ Rearrangement and Substitution Parameters Enables Accurate B Cell Receptor Sequence Annotation. PLoS Comput. Biol., 12, 1–25.

Safonova, Y. et al. (2015) IgSimulator: a versatile immunosequencing simulator. Bioinformatics, 31, 3213–3215.

Smakaj, E. et al. (2019) Benchmarking immunoinformatic tools for the analysis of antibody repertoire sequences. Bioinformatics, 36, 1731–1739.

Weber, C.R. et al. (2020) ImmuneSIM: Tunable multi-feature simulation of B- And T-cell receptor repertoires for immunoinformatics benchmarking. Bioinformatics, 36, 3594–3596.

Yaari, G. and Kleinstein, S.H. (2015) Practical guidelines for B-cell receptor repertoire sequencing analysis. Genome Med., 7, 121.

Yang, X. et al. (2021) Novel Allele Detection Tool Benchmark and Application With Antibody Repertoire Sequencing Dataset. Front. Immunol., 12, 739179.

Ye, J. et al. (2013) IgBLAST: an immunoglobulin variable domain sequence analysis tool. Nucleic Acids Res., 41, W34–W40.

Yermanos, A. et al. (2017) Comparison of methods for phylogenetic B-cell lineage inference using time-resolved antibody repertoire simulations (AbSim). Bioinformatics, 33, 3938–3946.

Zheng, B. et al. (2022) B-cell receptor repertoire sequencing: Deeper digging into the mechanisms and clinical aspects of immune-mediated diseases. iScience, 25, 105002.

