## Supplementary Methods for "AIRRSHIP: simulating human B cell receptor repertoire sequences"

### Reference datasets

The datasets used to generate the reference files included with AIRRSHIP are detailed in Supplementary Table 1. The source of data for each reference file is listed in Supplementary Table 2. Datasets in Supplementary Table 3 were used when comparing simulated repertoires to experimental. All reference and comparison datasets consist of human BCR sequences and only heavy chain sequences were analysed.

Raw sequence data were processed using pRESTO (Vander Heiden *et al.*, 2014) and Change-O (Gupta *et al.*, 2015) from the Immcantation analysis suite, with VDJ assignment carried out using IgBLAST v1.18.0 (Ye *et al.*, 2013). Pre-processed data were converted back into FASTA format using Change-O, and VDJ assignment was repeated using IgBLAST v1.18.0 to ensure consistency between datasets.

All reference data can also be replaced by the user and details are given in the documentation of required formats.

### Sequence generation process

**Locus creation –** AIRRSHIP takes IMGT FASTA files of VDJ alleles as input. These files are used to populate artificial heavy chain loci, each including single alleles representing every V, D and J gene. If more than two alleles exist for that gene, the allele included on each chromosome may differ (resembling heterozygous carriage) and the user can control the proportion of genes for which this is desired. By default, this means no one repertoire will contain sequences formed from more than two alleles of a single gene. However, the user can override this requirement and all alleles present in the input dataset will be used.

**VDJ usage –** To generate the initial recombined sequence, VDJ alleles are chosen based on gene usage distributions established from published datasets produced by 5' RACE amplification. Gene segments are recombined from the same synthetic loci only. Users may also specify VDJ usage to be even per gene or per gene family.

**Trimming –** The 3' end of the V gene, 5' and 3' end of the D gene and 5' end of the J gene are then trimmed. For each gene end, the number of nucleotides to be removed is sampled from distributions of trimming lengths for each IMGT gene family. To maintain sequence productivity, trimming lengths that would extend past the C - 104 anchor in the V gene or the W/F - 118 anchor in the J gene, or that would result in complete deletion of the D gene, are resampled. Users can also choose to not trim any or all of the gene ends.

**NP nucleotide addition –** AIRRSHIP does not distinguish between N and P nucleotides when insertions are modelled at the VD (NP1) and DJ (NP2) junctions. For each insertion, the number of nucleotides to be added is sampled from distributions of NP1 or NP2 lengths. The first nucleotide to be inserted is then randomly selected with probabilities for each base determined from public data for NP1 and NP2. Addition of further nucleotides follows a position dependent Markov process. For each position in the NP region, the next base is chosen with likelihood determined by a transition matrix that considers the current position in the sequence and the nucleotide base present at that position. When simulating non-mutated sequences and hypermutated sequences, a transition matrix determined from IgD/IgM sequences and from IgA/IgG sequences respectively, is used. This compensates for the inability to determine which positions within an NP region have been hypermutated. NP addition can be controlled with additions at either or both junctions optional.

**Somatic hypermutation –** Somatic hypermutation (SHM) is replicated at both the per sequence and per nucleotide position level. For each sequence, the overall mutation frequency is chosen from a distribution established from published datasets produced by 5’ RACE amplification methods. SHM rates can also be specified to be flat, such that every sequence has the same mutation frequency or number of mutations. Each 5mer within the sequence, excluding those where the centre base is an NP nucleotide, is then randomly iterated over. A new base is chosen to replace the centre base of this 5mer, sampled from a set of distributions that give the frequency of each nucleotide occurring at the centre position of each unique kmer for each sequence region (e.g. FWR1, CDR1). If the base chosen differs from the germline base, a mutation is introduced at that position and the process repeats until the number of mutations required to give the desired mutation frequency is reached. There is also the option for per position mutation to be context independent and the decision to mutate each base made entirely randomly.

**Productivity –** AIRRSHIP aims to produce only productive BCR sequences, defined here as those with no stop codons, the correct junction anchor residues (C-104 and W/F-118), and in-frame V and J segments. Alleles which do not have the correct anchor residues are excluded when processing the input data. Checks for productivity occur following trimming and NP addition. As certain VDJ combinations may be more likely to result in non-productive rearrangements, multiple attempts are made at trimming and nucleotide insertion using the same allele set to maintain VDJ usage distributions. SHM also commonly renders sequences non-productive, so multiple attempts at introducing hypermutation at the same rate will be made to ensure the per-sequence hypermutation reflects published datasets.

**Comparisons between simulated repertoires**

Four repertoires with at least 12,000 unique rearrangements, with and without SHM, were generated using each of the tools to be compared – AIRRSHIP (v0.1.2), immuneSIM (v0.8.7), IMPlAntS and partis (v0.15.0). Simulations were run as recommended in tool documentation and in such a way that required no additional reference data to be provided. We found some required parameters files were missing from IMPlAntS, so IMGT allele reference FASTA files were downloaded, and existing substitution model files duplicated to replace those absent for IGHV4 and IGHV5 genes.

VDJ assignment was carried out with IgBLAST, and the output processed into AIRR format TSV files using Change-O. We compared IgBLAST assignments from these sequences, rather than ground-truth output, so that any biases introduced by IgBLAST processing of the experimental repertoires would also be present in the simulated repertoires.

Output files for experimental and simulated repertoires were then further formatted for use with sumrep. Only sequences marked as productive and with a junction containing the correct anchor residues were kept. Where VDJ assignment was ambiguous, the first call was used for that sequence. Sequences were filtered to either IgD & IgM or IgA & IgG and where the repertoire size exceeded 10,000, samples were subset to this number of sequences. For partis specifically, one sequence was randomly chosen from each clone before subsampling. Deletion lengths for each gene end were calculated, the NP regions (VD/DJ insertions) were identified and a germline alignment with the D gene masked was produced.

Sumrep (Olson *et al.*, 2019) was installed from https://github.com/catsutherland/sumrep which contains a small change in the getMarkovMatrix function that ensures the transition matrix is correctly calculated. Summary measures were calculated for every repertoire and repertoires from each simulation tool were compared pairwise to each of the experimental repertoires and the divergence between them calculated. For VDJ gene usage, comparisons were made only to the 5’ RACE experimental datasets due to possible biases introduced by multiplex primer methods. This process was carried out for non-mutated simulated datasets in comparison with IgD/IgM experimental sequences, and for mutated simulated datasets in comparison with IgA/IgG experimental sequences.

To compare per position mutation frequency, the average mutation frequency of each position (number of times sequence position mutated divided by number of times sequence position present) across every IgA/IgG sequence in the 5’ RACE comparison datasets was calculated. The four repertoires generated from each simulated tool were condensed and the average per position mutation frequency compared with the experimental data. Only positions that occurred in at least 1% of sequences were included. The Pearson correlation coefficient was calculated using the SciPy stats package (Virtanen *et al.*, 2020).

**Runtime comparisons**

Each simulation tool was used to generate 10 repertoires, with and without mutation, of 10,000 VDJ recombinations. The shell time command, or the system.time command in R were used to calculate run time. All tools were run on an Intel (R) Xeon (R) CPU with 8 cores and 188Gb RAM. Eight processes were specified when running partis.

**AIRRSHIP VDJ assignment benchmarking example**

Three repertoires, each of 100,000 sequences were generated using AIRRSHIP. One with default settings, one using the --no-trim option and one with the -no_trim_d3 and --no_trim_d5 options. These repertoires were then analysed using both IMGT/HighV-Quest through the online portal, and IgBLAST v1.18.0 which was installed locally and run using the Change-O wrapper. The percentage of sequences where the length of the identified NP1 nucleotide sequence matched that known to be inserted by AIRRSHIP was calculated.

**Supplementary Table 1:** Experimental datasets used to generate reference files present in AIRRSHIP package. All samples were described as healthy or control individuals, and only heavy chain sequences were included.

| **Reference** | **Number samples** | **Downloaded data** | **Method** | **Comments** |
|---|---|---|---|---|
| Cowan, unpublished data | 92 | NA | Multiplex primers | Healthy individuals of Ghanaian descent. |
| Ghraichy *et al.*, 2020 | 42 | Pre-processed https://zenodo.org/record/3585046#.Y2upy4LP2JF | Multiplex primers | Only data from individuals aged over 9 years of age were included. |
| Gidoni *et al.*, 2019 | 48 | Raw sequence - ENA, accession PRJEB26509 | 5' RACE | Only healthy controls included. |
| Waltari *et al.*, 2018 | 14 | Raw sequence - SRA, accession PRJNA393446 | 5'RACE | NA |
| Yang *et al.*, 2021 | 184 | Raw sequence – SRA, accession PRJNA564936 | 5'RACE | Only data from individuals with at least 1000 unique sequences were included. |

**Supplementary Table 2:** Source of data and purpose of each file in the built-in AIRRSHIP reference data.

| **Reference file** | **Purpose** | **Source** |
|---|---|---|
| imgt_human_IGH[V\|D\|J].fasta | VDJ allele nucleotide sequences | IMGT reference directory (Lefranc and Lefranc, 2001) |
| IGH[V\|D\|J]_usage_gene.csv | VDJ usage | Waltari *et al.*, 2018; Gidoni *et al.*, 2019; Yang *et al.*, 2021 |
| [V\|D\|J]_family_trimming_proportions.csv | Gene end trimming | Cowan, unpublished data; Waltari *et al.*, 2018; Gidoni *et al.*, 2019; Ghraichy *et al.*, 2020; Yang *et al.*, 2021 |
| np[1\|2]_first_base_probs.csv | First base usage for NP regions | Cowan, unpublished data; Waltari *et al.*, 2018; Gidoni *et al.*, 2019; Ghraichy *et al.*, 2020; Yang *et al.*, 2021 |
| np[1\|2]_transition_probs_per_position_ig[dm\|ag].csv | Transition matrix for NP insertions | Cowan, unpublished data; Waltari *et al.*, 2018; Gidoni *et al.*, 2019; Ghraichy *et al.*, 2020; Yang *et al.*, 2021 |
| mut_freq_per_seq_per_family.csv | Mutation frequency per sequence distribution | Waltari *et al.*, 2018; Gidoni *et al.*, 2019; Yang *et al.*, 2021 |

| | | |
|---|---|---|
| [region]_kmer_base_usage.csv | Mutation likelihoods for each unique 5mer | Waltari *et al.*, 2018; Gidoni *et al.*, 2019; Yang *et al.*, 2021 |

**Supplementary Table 3:** Experimental datasets used as comparators. All samples were described as healthy or control individuals, and only heavy chain sequences were included.

| Reference | Number samples | Sumrep samples | Data download | Method | Comments |
|---|---|---|---|---|---|
| Briney *et al.*, 2019 | 10 | 327059, 326651, 326713, 326797 | Raw sequence - SRA, accession PRJNA406949 | Multiplex primers | One technical replicate of the first biological replicate from each individual was used (files >2M reads were downsampled). |
| Cowan *et al.*, 2020 | 16 | B1639, B1413, B6844, B4077 | Raw sequence - ENA, accession PRJNA561156 | Multiplex primers | NA |
| Gupta *et al.*, 2017 | 3 | B, MC, V | Pre-processed https://zenodo.org/record/821659#.YPBKLm7TXmF | 5' RACE | Only sequence data from timepoints prior to vaccination were used. |
| Rubelt *et al.*, 2016 | 10 | TW04A, TW04B, TW02A, TW05A | Pre-processed - VDJServer | 5' RACE | NA |
| Vander Heiden *et al.*, 2017 | 4 | HD07, HD09, HD10, HD13 | Raw sequence - SRA, accession PRJNA338795 | 5'RACE | NA |

**References**


Briney,B. *et al.* (2019) Commonality despite exceptional diversity in the baseline human antibody repertoire. *Nature*, **566**, 393–397.

Cowan,G.J.M. *et al.* (2020) In Human Autoimmunity, a Substantial Component of the B Cell Repertoire Consists of Polyclonal, Barely Mutated IgG+ve B Cells . *Front. Immunol.* , **11**, 395.

Ghraichy,M. *et al.* (2020) Maturation of the Human Immunoglobulin Heavy Chain Repertoire With Age. *Front. Immunol.*, **11**, 1734.

Gidoni,M. *et al.* (2019) Mosaic deletion patterns of the human antibody heavy chain gene locus shown by Bayesian haplotyping. *Nat. Commun.*, **10**, 628.

Gupta,N.T. *et al.* (2015) Change-O: a toolkit for analyzing large-scale B cell immunoglobulin repertoire sequencing data. *Bioinformatics*, **31**, 3356–3358.

Gupta,N.T. *et al.* (2017) Hierarchical Clustering Can Identify B Cell Clones with High Confidence in Ig Repertoire Sequencing Data. *J. Immunol.*, **198**, 2489–2499.

Vander Heiden,J.A. *et al.* (2017) Dysregulation of B Cell Repertoire Formation in Myasthenia Gravis Patients Revealed through Deep Sequencing. *J. Immunol.*, **198**, 1460 LP – 1473.

Vander Heiden,J.A. *et al.* (2014) pRESTO: a toolkit for processing high-throughput sequencing raw reads of lymphocyte receptor repertoires. *Bioinformatics*, **30**, 1930–1932.

Lefranc,M.-P. and Lefranc,G. (2001) The Immunoglobulin FactsBook Academic Press, London.

Olson,B.J. *et al.* (2019) sumrep: A Summary Statistic Framework for Immune Receptor Repertoire Comparison and Model Validation. *Front. Immunol.*, **10**, 2533.

Rubelt,F. *et al.* (2016) Individual heritable differences result in unique cell lymphocyte receptor repertoires of naïve and antigen-experienced cells. *Nat. Commun.*, **7**, 11112.

Virtanen,P. *et al.* (2020) SciPy 1.0: fundamental algorithms for scientific computing in Python. *Nat. Methods 2020 173*, **17**, 261–272.

Waltari,E. *et al.* (2018) 5′ Rapid Amplification of cDNA Ends and Illumina MiSeq Reveals B Cell Receptor Features in Healthy Adults, Adults With Chronic HIV-1 Infection, Cord Blood, and Humanized Mice. *Front. Immunol.*, **9**, 628.

Yang,X. *et al.* (2021) Large-scale analysis of 2,152 Ig-seq datasets reveals key features of B cell biology and the antibody repertoire. *Cell Rep.*, **35**, 109110.

Ye,J. *et al.* (2013) IgBLAST: an immunoglobulin variable domain sequence analysis tool. *Nucleic Acids Res.*, **41**, W34–W40.
