## Supplementary Figures for "AIRRSHIP: simulating human B cell receptor repertoire sequences"

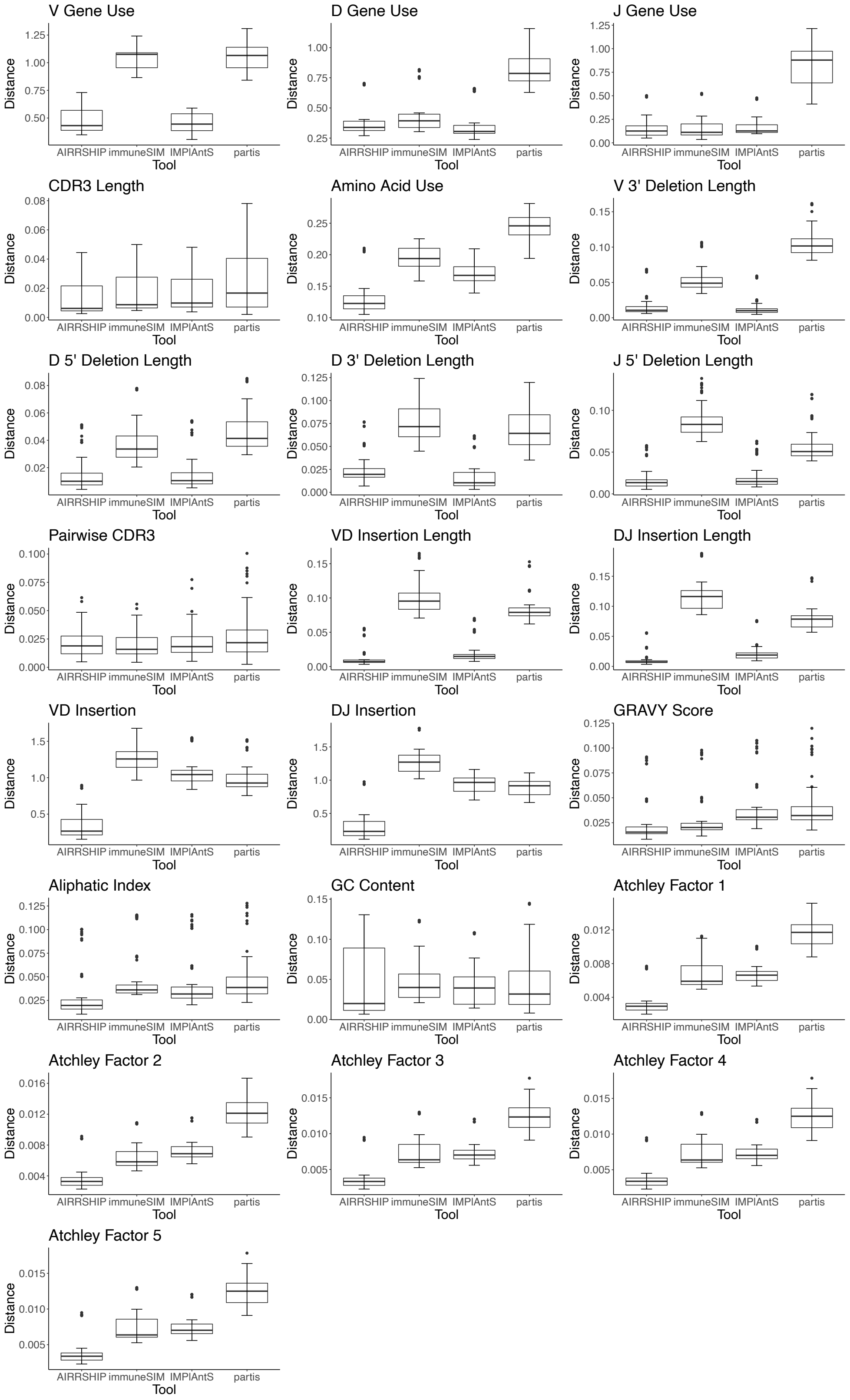


**Supplementary Figure 1**: Plots of summary distance values calculated using sumrep – unmutated simulations and IgD/IgM comparisons. For each tool, four simulated repertoires were each compared pairwise to a set of experimental repertoires.

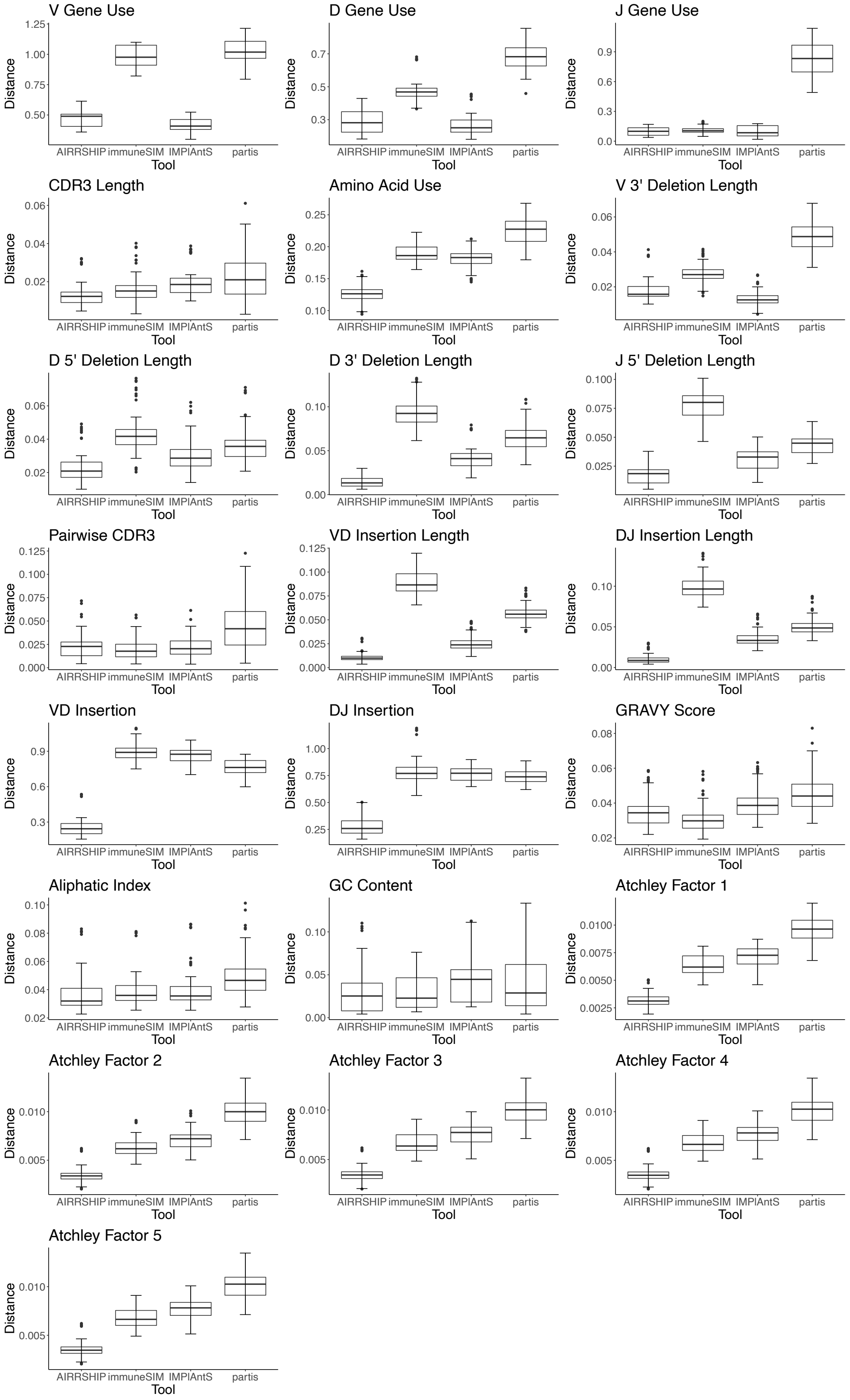


**Supplementary Figure 2**: Plots of summary distance values calculated using sumrep – mutated simulations and IgA/IgG comparisons. For each tool, four simulated repertoires were each compared pairwise to a set of experimental repertoires.

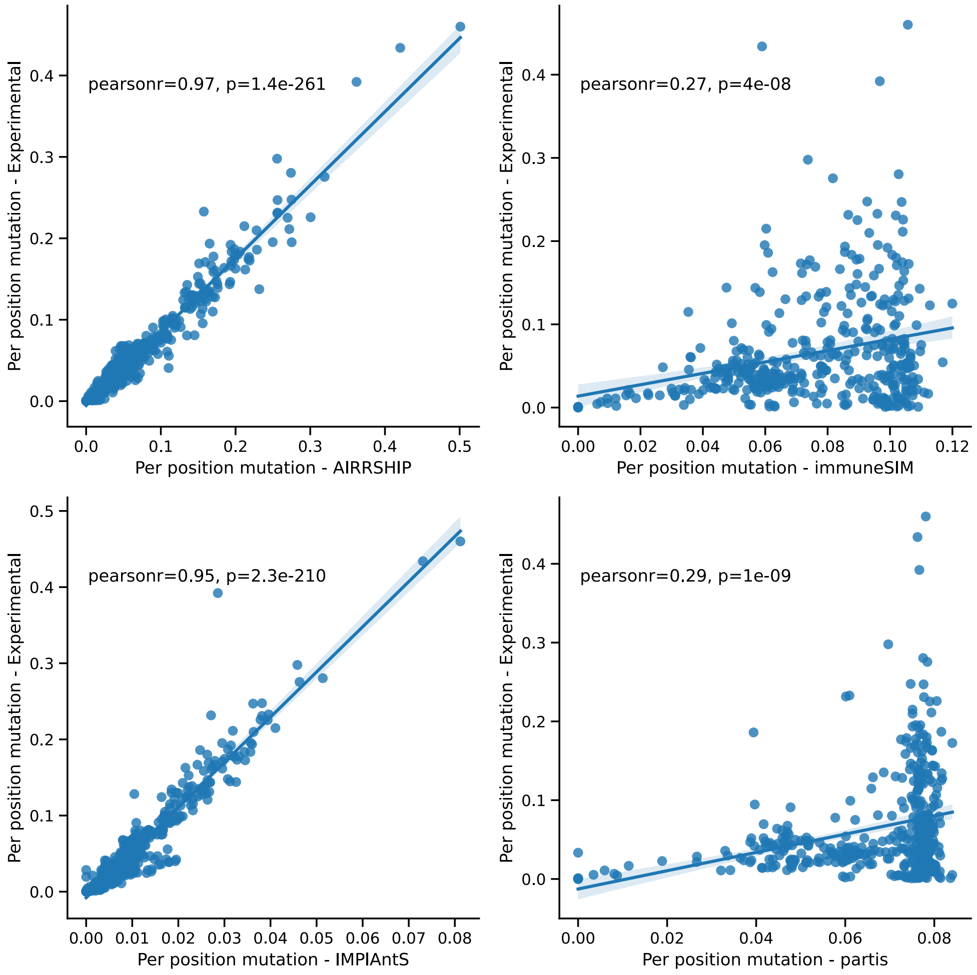


**Supplementary Figure 3**: Correlation of per position mutation across sequences between experimental and simulated sequences. For each tool, the four simulated repertoires were combined and compared to the entire set of experimental sequences. Each point represents a single position in the IMGT unique numbering.

**Supplementary Table 4**: Runtimes for creation of 10,000 unique VDJ rearrangements, with and without somatic hypermutation. Runtime was measured across ten simulations using either the shell time command or the R system.time() function and is displayed as mean (range). Partis was run using eight processes, all other tools were run as default as no thread specification was possible.

| **Tool** | **Mean real time (seconds)** | **Mean user time (seconds)** | **Mean real time – SHM (seconds)** | **Mean user time – SHM (seconds)** |
|---|---|---|---|---|
| AIRRSHIP | 7.7 (7.5 – 7.8) | 7.6 (7.5 – 7.8) | 65.5 (62.4 – 68.5) | 64.5 (62.2 – 67.3) |
| immuneSIM | 606.4 (598.4 – 618.1) | 606.1 (598.1 – 617.8) | 3561.4 (3494.1 – 3622.8) | 3559.2 (3492.5 – 3621.2) |
| IMPlAntS | 30.4 (30.2 – 30.5) | 20.5 (30.3 – 30.7) | 135.8 (133.7 – 137.8) | 135.6 (133.5 – 137.5) |
| partis | 100.4 (99.9 – 101.1) | 512.0 (510.5 – 513.7) | 133.2 (131.5 – 135.9) | 691.1 (685.9 – 703.7) |

**Supplementary Table 5:** Runtimes for creation of 10,000; 100,000 and 1,000,000 unique VDJ rearrangements, with and without somatic hypermutation. Runtime was measured across ten or five simulations (10,000 and 100,000/1,000,000 sequences, respectively) using the shell time command and is displayed as mean (range).

| **Number of sequences** | **Mean real time (seconds)** | **Mean real time – SHM (seconds)** |
|---|---|---|
| 10,000 | 7.7 (7.5 – 7.8) | 65.5 (62.4 – 68.5) |
| 100,000 | 77.0 (75.5 – 78.0) | 658.7 (654.4 – 661.6) |
| 1,000,000 | 745.5 (737.6 – 755.5) | 6495.8 (6402.8 - 6644.5) |

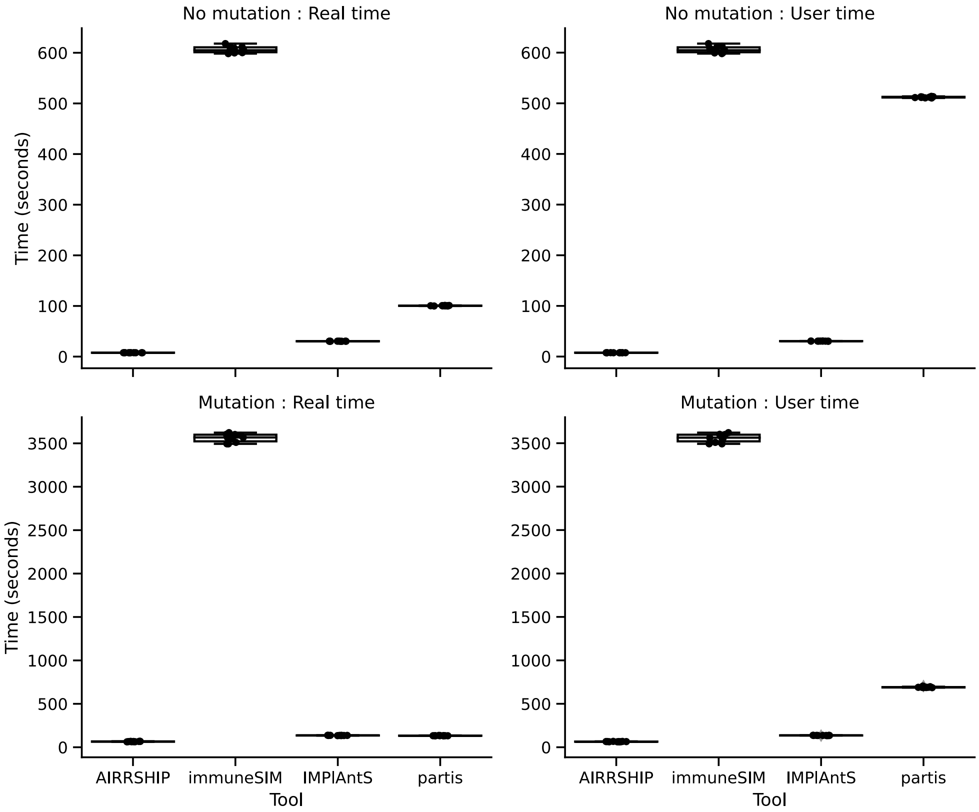


**Supplementary Figure 4:** Runtimes for simulation of 10,000 VDJ recombinations for each simulation tool, with and without the introduction of somatic hypermutation. Real time is equivalent to wall clock time, user time indicates the CPU time involved in the process. These differ only significantly for partis which can use multiple threads.

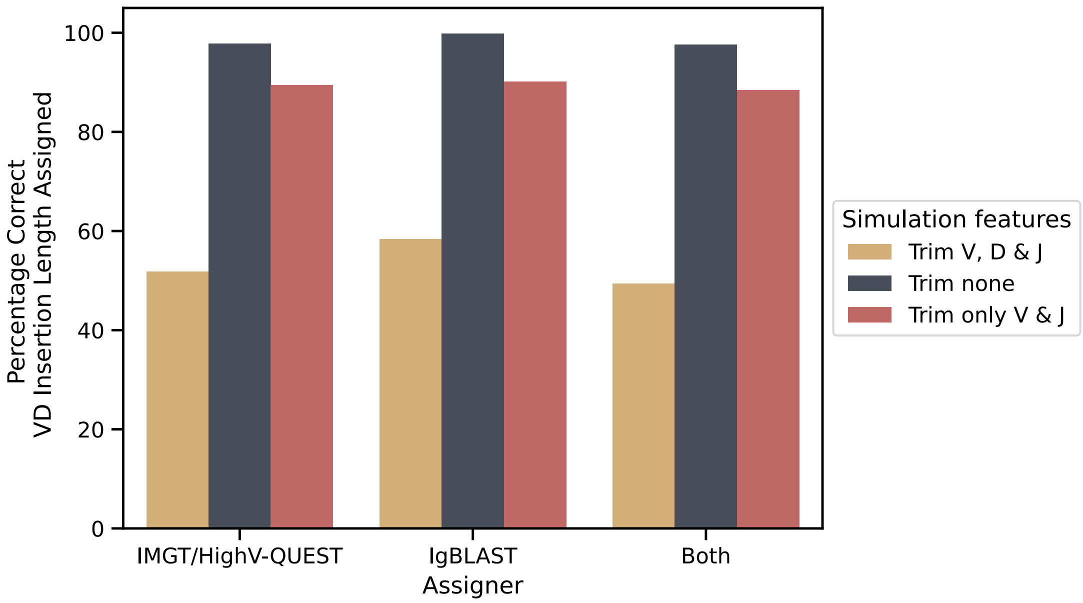


**Supplementary Figure 5**: Comparison of percentage VD insertion lengths correctly assigned by IgBLAST and IMGT/HighV-QUEST when run on repertoires simulated by AIRRSHIP using different trimming parameters.
